# CLUEY enables knowledge-guided clustering and cell type detection from single-cell omics data

**DOI:** 10.1101/2024.11.14.623697

**Authors:** Daniel Kim, Carissa Chen, Lijia Yu, Jean Yee Hwa Yang, Pengyi Yang

**Affiliations:** Computational Systems Biology Unit, Children’s Medical Research Institute, Faculty of Medicine and Health, University of Sydney, NSW 2145, Australia; School of Mathematics and Statistics, Faculty of Science, University of Sydney, NSW 2006, NSW, Australia; Sydney Precision Data Science Centre, University of Sydney, Sydney, NSW 2006, Australia; Charles Perkins Centre, University of Sydney, Sydney, NSW 2006, Australia

## Abstract

Clustering is a fundamental task in single-cell omics data analysis and can significantly impact downstream analyses and biological interpretations. The standard approach involves grouping cells based on their gene expression profiles, followed by annotating each cluster to a cell type using marker genes. However, the number of cell types detected by different clustering methods can vary substantially due to several factors, including the dimension reduction method used and the choice of parameters of the chosen clustering algorithm. These discrepancies can lead to subjective interpretations in downstream analyses, particularly in manual cell type annotation. To address these challenges, we propose CLUEY, a knowledge-guided framework for cell type detection and clustering of single-cell omics data. CLUEY integrates prior biological knowledge into the clustering process, providing guidance on the optimal number of clusters and enhancing the interpretability of results. We apply CLUEY to both unimodal (e.g. scRNA-seq, scATAC-seq) and multimodal datasets (e.g. CITE-seq, SHARE-seq) and demonstrate its effectiveness in providing biologically meaningful clustering outcomes. These results highlight CLUEY on providing the much-needed guidance in clustering analyses of single-cell omics data. CLUEY package is available from https://github.com/SydneyBioX/CLUEY.

## Introduction

Advancements in single-cell omics technologies, especially their progression towards multimodality, have transformed our ability to profile global molecular attributes at single-cell resolution. Clustering is a critical task in single-cell omics data analysis, serving as a foundational step for detecting putative cell types before downstream analyses such as differential expressed gene identification and cell type annotation (Heumos et al., 2023). The standard approach involves first grouping cells based on their molecular attributes, such as gene expression profiles, and then manually annotating each cluster using distinct marker genes identified through differential expression analysis. The number and the quality of the clusters generated by clustering algorithms can have a significant impact on downstream analyses and their biological interpretation (Butler et al., 2018). Yet, for a given dataset, substantial differences in clustering results are often observed (Duò et al., 2020; Freytag et al., 2018). For instance, different clustering methods applied to the same dataset can produce results with substantially different number of clusters (Yu et al., 2022). Additionally, factors such as the choices of dimension reduction techniques, similarity metrics (Kim et al., 2019), and clustering algorithm parameters further contribute to the variability in clustering results. These discrepancies often lead to subjective interpretations, complicating downstream analyses such as cell type detection and annotation.

To address these challenges, methods such as SC3 attempt to stabilise clustering results by combining multiple clustering methods through a consensus approach (Kiselev et al., 2017), whereas methods such as Seurat indirectly achieve clustering consensus by integrating gene expression data with additional data modalities (e.g. surface protein abundance) (Hao et al., 2021). Alternatively, various methods have been developed to actively estimate the number of cell types in the data using inherent data structures such as inter- and intra-cluster similarities, eigenvector-based metrics, and clustering stability metrics (Yu et al., 2022). Despite these improvements, most clustering algorithms for single-cell data rarely incorporate prior biological knowledge during the clustering process. This omission can result in clusters that do not align with biologically meaningful cell types. Incorporating known biological information, such as known cell type markers, could improve the interpretability of the clustering results and guide the algorithm towards more biologically relevant clusters. This can reduce the risk of generating clusters that are artifacts of the algorithm or driven primarily by statistical patterns rather than biological signals, ultimately improving downstream analyses such as differential expression analysis and cell type detection and annotation.

Here, we propose CLUEY, a knowledge-guided framework for estimating the number of cell types and clustering of both unimodal and multimodal single-cell omics data. Building on concepts from our previous work on optimising clustering with prior biological knowledge (Yang et al., 2015), CLUEY identifies and leverages cell identity genes (Kim et al., 2021) associated with a wide range of cell types from large-scale single-cell atlases to construct knowledgebases and subsequently uses them to inform the clustering of query data, determining the most representative number of clusters. This is achieved by recursively clustering until an optimal number of cell types is detected based on the knowledgebase. By integrating prior knowledge, CLUEY performs clustering in an interpretable and biologically informed manner. Additionally, CLUEY provides interpretable enrichment statistics associated with the optimal clustering, facilitating further downstream processing and exploration. We evaluate the performance of CLUEY on both unimodal and multimodal single-cell omics data and benchmark it against popular state-of-the-art methods. Our results demonstrate CLUEY’s ability and robustness in estimating the number of cell types across a range of datasets with various data characteristics, showcase its utility in annotating these cell types using the knowledgebase, and highlight CLUEY as a novel approach for clustering optimisation and cell type annotation in single-cell omics data analysis.

## Methods

### Data Collection

#### Mouse and Human Cell Atlases

The Tabula Muris Atlas (a.k.a. Mouse Cell Atlas [MCA]) (Schaum et al., 2018) and Tabula Sapiens Atlas (a.k.a. Human Cell Atlas [HCA]) (The Tabula Sapiens Consortium* et al., 2022) each comprise two versions: one generated using Fluorescence-Activated Cell Sorting (FACS) and Smart-Seq2 sequencing protocols, and the other using 10x Genomics technology. Lowly expressed genes with less than a total of 10 counts and cells with less than a total of 200 counts were removed from both datasets. Gene expression values were then normalized and log-transformed using the LogNormalize function in Scater (McCarthy et al., 2017). Additionally, cell types with fewer than 20 cells were excluded from both atlases in the evaluation.

The MCA (FACS) includes 23,433 genes, 44,779 cells, and encompasses 81 cell types from 20 organs. The MCA (10X) retains the same number of genes but consists of 54,865 cells and includes 55 cell types from 12 organs. Similarly, the HCA (FACS) dataset contains 58,870 genes, 27,051 cells, and encompasses 133 cell types from 24 organs. In contrast, the HCA (10X) dataset, also composed of 32,911 genes, includes 181,775 cells and 132 cell types from 24 organs.

#### TEA-seq dataset

The processed data of TEA-seq data from measuring PBMC were downloaded from the NCBI Gene Expression Omnibus (GEO) under the accession number GSM5123954, with raw RNA expression and peak accessibility (ATAC) measured for the same cells. We summarized the matrix of ATAC from peak level to gene activity scores using the ‘CreateGeneActivityMatrix’ function in the Seurat v3 package (Hao et al., 2021). Genes with fewer than 1% quantifications, including those with zero counts, across cells in each of the two modalities were removed. After preprocessing, we retained 9,661 genes in the RNA modality and 16,434 chromatin accessible regions in the ATAC modality, with 6,507 cells and nine cell types in both modalities.

#### CITE-seq (E-MTAB-10026) dataset

This CITE-seq dataset measures PBMC from healthy individuals and from COVID-19 patients (Stephenson et al., 2021). Only the data from healthy individuals were used in this study. The raw matrices of RNA and ADT and the annotation of cells to their respective cell types from the original study were downloaded from the EMBL-EBI ArrayExpress database under the accession number E-MTAB-10026. RNA and ADT in this dataset were filtered by removing those that expressed in less than 1% of the cells, including those with zero counts, and cell types were filtered by removing those that have less than 10 cells. After filtering, there are 10,665 genes and 192 surface proteins in the RNA and ADT modalities, respectively, including 30,271 cells from 16 cell types.

### CLUEY algorithm

#### Overview

The CLUEY algorithm is composed of three components, excluding the generation of the knowledgebase (Figure 1). First, given a query dataset, the dimensions of the data are reduced using a multimodal autoencoder (**dimensionality reduction**). The second component involves the initial estimation of the optimal number of cell types in a uni- or multimodal query dataset, using a knowledgebase. The third and final step is **recursive clustering**, which recursively clusters the query data to capture finer cell type populations and more thoroughly determine the final prediction of the optimal number of cell types in the data. CLUEY requires gene sets of known cell types to inform the clustering process. We generate sets of the top differentially stable genes for each cell type in a given reference gene expression matrix using Cepo (Kim et al., 2021). The expression profiles of the differentially stable genes are then averaged across the cell types in the knowledgebase to generate their respective pseudo-bulk profiles. For broader usability, we have also implemented the option for creating pseudo-bulk profiles of differentially expressed genes from a knowledgebase that have a Bonferroni corrected p-value < 0.05 for each cell type in the knowledgebase using Seurat. It is also possible to provide a set of any manually determined gene sets and their corresponding averaged expression values as the knowledgebase to CLUEY. Below we give a detailed description of each of the three components of CLUEY’s algorithm.

**Figure 1.**
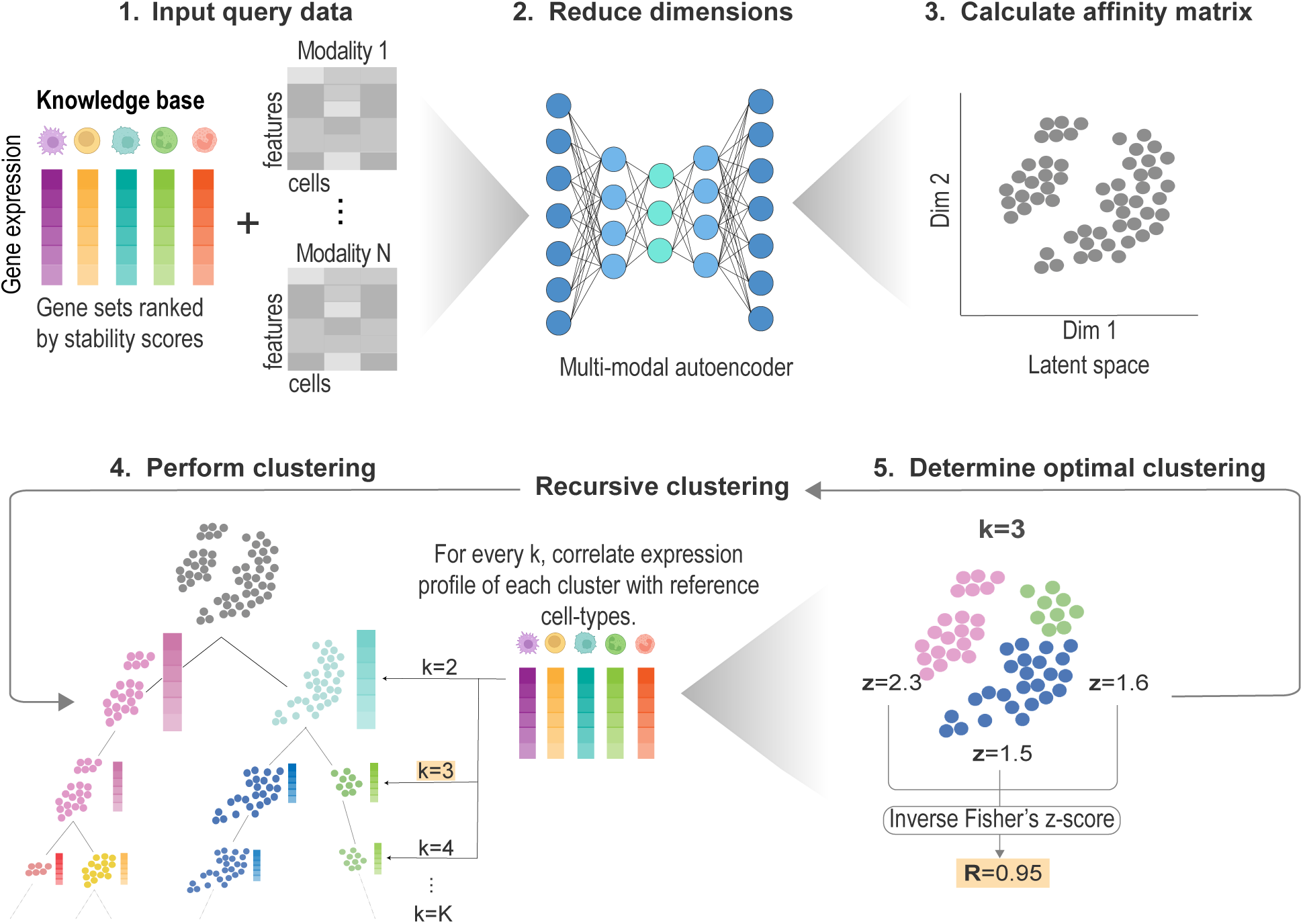
Overview of the CLUEY Algorithm. Steps 1-3: CLUEY builds a knowledgebase of cell types and projects a uni- or multimodal query dataset to a latent space using an autoencoder before calculating an affinity matrix. Step 4: CLUEY performs clustering using the spectral algorithm and the knowledgebase to determine the initial optimal number of clusters. The expression profiles of each putative cluster are then averaged and correlated to the gene sets in the knowledgebase. The most correlated cell type in the knowledgebase is used to determine biologically-informed clustering. Step 5: The correlation for each cluster is transformed to a z-score using Fisher’s method and averaged using the inverse of Fisher’s method. This is the clustering score for the entire grouping. Steps 4 and 5 are repeated recursively for each initial cluster identified in step 4, until an optimal number of clusters is reached.

#### Component 1 - Dimensionality reduction

First, we select the top 5% highly variable genes (HVGs) for input into a simple multi-layered autoencoder, inspired by Matilda (Liu et al., 2023). The resulting 10 encodings, which can be tuned, are extracted and used to construct an affinity matrix, by calculating the Euclidean distance between query cells. For training using multimodal data, each modality is passed through an encoder and concatenated into a single latent space. In the decoding phase, the merged branch is split into modality-specific decoders. We then extract the shared latent space for clustering. Hyperparameters such as the learning rate, number of hidden layers, and encodings can be specified by the user and adjusted according to the needs of the query dataset.

#### Component 2 - Clustering

To determine the optimal number of clusters in a given query dataset, we cluster the data from *k=2* to a user-defined *K* number of clusters (*K=20* as the default). For each *K*, pseudo-bulk gene expression profiles are generated for each predicted cluster and the intersecting genes between the cluster pseudo-bulk profile and top 10% of genes (default parameter) for each reference cell type’s genes are correlated using Spearman’s correlation. We weight each pairwise correlation to ensure that we account for the number of genes used when correlating the pseudo-bulk profiles. For a given number of cluster *k*, the weighted correlation coefficients are then transformed using Fisher’s Z transformation (*z_ij_*) for a given cluster *i* and a cell type j is written as

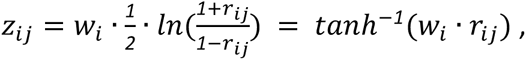

where *w* represents the weight for cluster *i* and *C* is the number of cell types in the knowledgebase and *r_ij_* is the correlation coefficient between the cluster *i* and cell type *j*.

We then average the z-scores for each grouping of clusters (*k=1, 2, 3, …K*) and use the inverse of Fisher’s Z transformation to get a correlation coefficient for each clustering. The optimal *Kth* grouping is determined by the maximum correlation coefficient *r*. Thus, we treat *r* as a measure to determine the optimal *K* clusters in a query dataset:

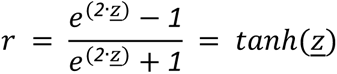

The initial prediction for the optimal number of clusters is simply the grouping with the largest correlation coefficient r. These clusters will then be re-clustered using CLUEY’s recursive clustering algorithm until no less than 30 cells are reached for each cluster. Importantly, the stopping condition can be specified such that the algorithm continues to recluster until no less than a minimum number of cells is specified by the user (default = 30 cells).

#### Component 3 - Recursive clustering

Importantly, CLUEY performs recursive clustering to capture any subpopulations that may have been missed by recursively reclustering the clusters identified in the previous step. This process is repeated until there are no less than 30 cells left for each subcluster, by default. However, we only reduce the dimensions once after the first level of reclustering and use this latent space for the subsequent steps. To minimize noise and account for the lower number of cells we take the top 5% HVGs and extract two dimensions. Thus, similar to bagging, we treat every initial cluster from the first round of clustering as a smaller query dataset. The final prediction for the optimal number of clusters is then returned, including the respective correlation coefficient and the closest matching reference cell types for interpretability.

### Settings for alternative methods

#### Seurat

We used Seurat (v5.0.2) to process and cluster the query data as outlined in the author’s vignette (https://satijalab.org/seurat) (Hao et al., 2021). First, the raw RNA count matrices were used as input and log-transformed using the ‘NormalizeData’ function in Seurat. We used the default setting and thus selected the top 2000 highly variable genes using the ‘FindVariableFeatures’ function before using the ‘runPCA’ function to reduce the dimensions of the data. We then used the ‘FindNeighbors’ and ‘FindClusters’ functions to determine the predicted clusters in the query dataset.

We also ran Seurat on multimodal data with similar steps taken as above. Here, we used the parameters of normalization.method = ‘CLR’ and margin = 2 in the in ‘NormalizeData’ function, as suggested in the author’s pipeline. PCA was performed using the ‘runPCA’ function and the function ‘FindMultiModalNeighbors’ integrates RNA and ADT/ATAC modalities using the PCA results. The joint visualization of RNA and ADT/ATAC were generated using the ‘wnn.umap’ function.

#### Monocle3

We used the latest version of Monocle3 (v1.3.7) with default parameter settings and followed the pipeline specified by the author (https://cole-trapnell-lab.github.io/monocle3) (Cao et al., 2019). Briefly, each dataset was initiated as a Monocle3’s main class ‘cell_data_set’, and then preprocessed before reducing the dimensions of the data using the ‘preprocess_cds’ and ‘reduce_dimension’ functions, respectively. Cells in the query data were then assigned predicted cluster membership using the ‘cluster_cells’ function.

#### SC3

We used SC3 (v1.32.0) (Kiselev et al., 2017) with default settings as outlined in the author’s tutorial (https://nbisweden.github.io/workshop-archive/workshop-scRNAseq/2018-05-21/labs/sc3_ilc.html). First, we estimated the initial number of cell types in the normalized and log-transformed query data using the ‘sc3_estimate_k’ function. The final predictions were extracted by calling the ‘sc3’ function using the initial estimate as a parameter.

#### CIDR

Cidr (v0.1.5) (Lin et al., 2017) with the default parameters was used on the normalized and log-transformed data to estimate the number of cell types in the query data (https://github.com/VCCRI/CIDR). The query data was converted to an scData object using the ‘scDataConstructor’ function. Dropout candidates and the dissimilarity matrix were calculated using the ‘determineDropoutCandidates’ and ‘scDissim’ functions, respectively, before reducing the dimensions of the data using ‘scPCA’. Finally, the number of predicted clusters was extracted after running ‘scCluster’.

#### RaceID

We used RaceID (v0.3.5) (Grün et al., 2015) to acquire the predicted number of clusters in the query data. The query data was first converted to an SCseq object before filtering the data using the ‘filterdata’ function. Distances between the cells were calculated using the ‘compdist’ function. We specified Pearson’s correlation as the distance metric (default) and extracted the predicted clusters using the ‘clustexp’ function with default parameters (https://github.com/dgrun/RaceID3_StemID2_package/blob/master/README.md).

#### SHARP

The raw counts were provided as input to SHARP (v1.1.0) (Wan et al., 2020). Clusters were then predicted using the ‘SHARP’ function, expression type was set to “count” and prep was set to true to preprocess the data and remove any non-expressed genes from the query data.

#### Spectrum

Spectrum (v1.1) was used with default parameters, where the maximum number of cell types that could be predicted was set to 30 (John et al., 2020). Cluster assignments were determined using the ‘Spectrum’ function and raw counts were provided as the query data.

### Performance Evaluation

We conducted comprehensive testing of CLUEY’s performance against seven methods across four distinct scenarios. Our evaluation focused on assessing CLUEY’s capability to accurately estimate the number of cell types in a query datasets and handle imbalance data. Additionally, we examined CLUEY’s clustering capabilities across four independent datasets, comprising two human and two mouse datasets.

#### Scenario one

we assessed CLUEY’s ability to annotate and estimate the number of cell types in a query dataset by subsampling the MCA (10X) and HCA (FACS). We excluded cell types with fewer than 200 cells and varied the number of cell types from five to 20 while keeping the number of cells for each cell type constant at 200. Thus, we generated 10 datasets for each specified number of cell types. This resulted in 40 datasets for each atlas.

#### Scenario two

we investigated the impact of imbalanced data by creating major and minor cell types. Ten cell types were randomly selected from each atlas and categorized into major and minor groups. We adjusted the cell counts within these groups to establish imbalance ratios of 2:1, 4:1, and 8:1 between the major and minor cell types. For instance, a 2:1 ratio indicates 200 cells in major cell types and 100 cells in minor cell types. This process was iterated 10 times for each ratio, yielding a total of 30 datasets.

#### Scenario three

we evaluated CLUEY on independent datasets, which included two human and two mouse datasets (Baron et al., 2016; Segerstolpe et al., 2016; Zeisel et al., 2015; Zilionis et al., 2019). To estimate variability, we randomly subsampled each of the four independent datasets to retain approximately 90% of the original number of cells and repeated this process ten times, resulting in 40 datasets for each species.

#### Scenario four

Finally, we evaluated CLUEY on multimodal data using the TEA-seq (Swanson et al., 2021) and CITE-seq (Stephenson et al., 2021) datasets. In the same manner as the third setting, we down-sampled the data to 90% of the original number of cells and repeated this 10 times to capture variability. This yielded a total of 20 datasets, 10 for each sequencing platform. To assess CLUEY’s ability to cluster a multimodal dataset using a knowledgebase generated uni-modal data, we clustered the datasets using the knowledgebase generated from the HCA (10x), which only contains gene expression values (RNA).

### Cell clustering evaluation metrics

To assess cluster concordance of CLUEY and the other tested methods, we used three evaluation metrics: Adjusted Rand Index (ARI), and Absolute Deviation as defined below:

Given a set S of n elements, and two groupings or vectors of cluster labels of these labels, namely *X* = {*X_1_*, *X_2_*, . . . , *X_n_*} and *Y* = {*Y_1_*, *Y_2_*, . . . , *Y_n_*}, the overlap between X and Y can be summarized in a contingency table [*n_ij_*] where each entry *n_ij_* denotes the number of elements in common between *X_i_* and *Y_j_*. Furthermore, *n_ij_* = |*X_i_* ∩ *Y_j_*|. Thus, the Adjusted Rand Index (ARI) is defined as:

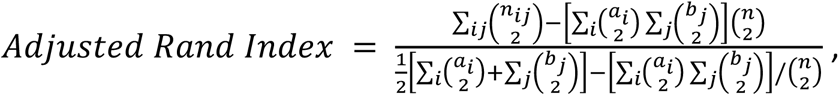

where *n_ij_*, *a_i_*, *b_j_* are values from the contingency table defined above.

To quantify the difference between the predicted number of clusters vs the true number of clusters, we took the absolute value of their difference as shown below:

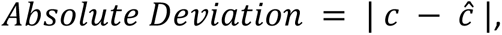

where c is the true number of cell types and *ĉ* is the predicted number of cell types.

## Results

### CLUEY enable cell clustering and the number of cell type estimation

We compared CLUEY’s performance against seven alternative methods using the Mouse Cell Atlas (MCA) and Human Cell Atlas (HCA) datasets. Our evaluation focused on two key areas, including the ability to estimate the true number of cell types (deviation) and the quality of cell clustering, where we assume a cluster corresponds to a cell type and quantified the concordance using ARI against the predefined cell type labels. First, we generated 40 balanced datasets for each atlas by subsampling the data and varying the number of cell types. The query datasets for MCA were sequenced using FACS, while those for HCA were sequenced using 10X Chromium. To assess CLUEY’s generalisability for cross-platform predictions, we generated knowledgebases for CLUEY from both the 10X and FACS-sequenced data and assessed its performance on FACS- and 10X- sequenced query data, respectively. Results from MCA data analyses demonstrated that CLUEY is highly competitive in estimating the number of cell types in the data (Figure 2A) and lead to highly concordant cell clustering compared to predefined cell type labels from MCA (Figure 2B). While its performance on estimating the number of cell types in HCA is less striking compared to other methods, the quality of the cell clustering as quantified by ARI remain high across HCA datasets (Figure 2C, D).

**Figure 2.**
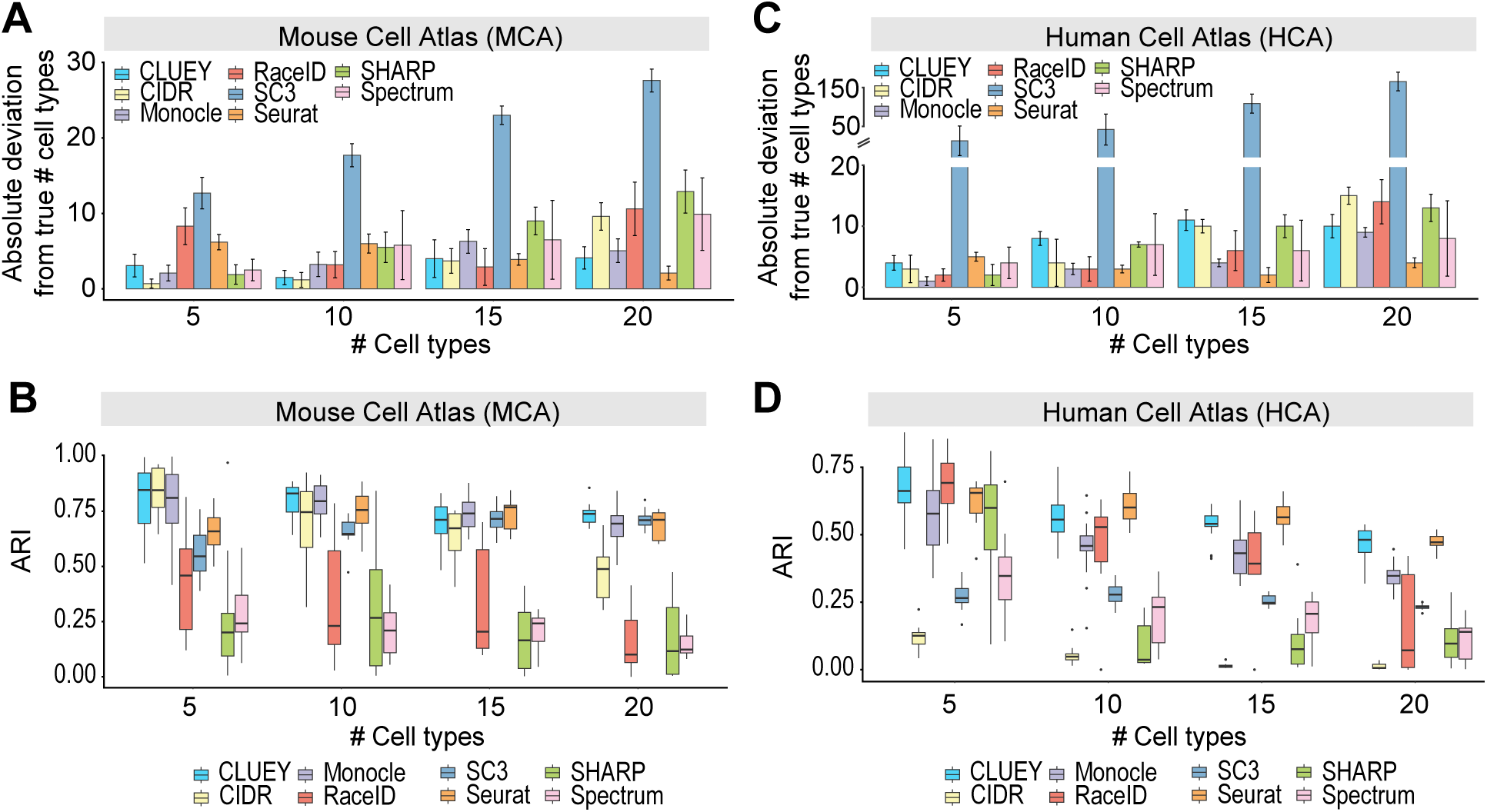
Benchmarking CLUEY on MCA and HCA datasets. **(A,C)** For each method, the absolute deviation between the estimated number of clusters and true number of cell types for (A) MCA, and (C) HCA. The number of cell types in the query datasets were increased from 5 to 10, 15, and then 20. **(B, D)** For each method, ARI quantification of cell clustering concordance with predefined cell type labels from for (B) MCA, and (D) HCA. Same as above, the number of cell types in the query datasets were increased from 5 to 10, 15, and then 20.

Next, we also tested CLUEY’s ability to correctly estimate the number of cell types in imbalanced query data. In comparison to alternative methods, CLUEY’s deviation from the true number of cell types remain low and stable across varying imbalance ratios (Figure 3A). This is largely mirrored by the quality of cell clustering, where CLUEY achieves best performance in most cases across all imbalance ratios (Figure 3B). Overall, these results demonstrate CLUEY’s utility in cell clustering and cell type detection.

**Figure 3.**
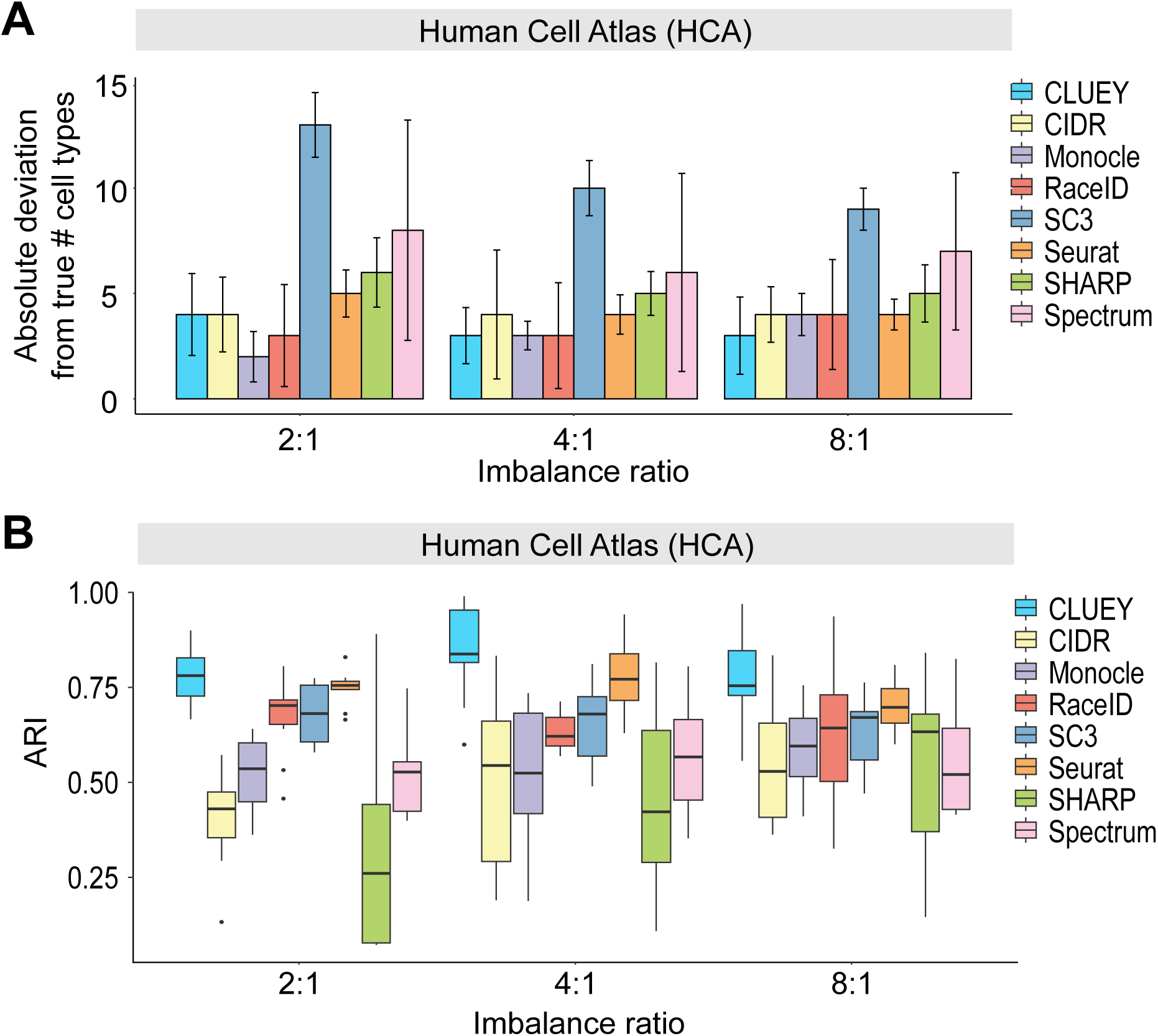
Evaluation of CLUEY and alternative methods on imbalanced data. **(A)** Absolute deviation on number of cell type estimation for each method on the HCA datasets. X-axis represents the imbalance ratio between the major and minor cell types. Thus, an imbalance ratio of 2:1 means there are 200 and 100 cells in the major and minor groups, respectively. Error bars represent two standard errors above the means. **(B)** ARI quantification of cell clustering for each method on the same HCA datasets as in (A).

### CLUEY demonstrates robust performance on knowledgebase generated from independent data sources

Single-cell RNA-seq datasets are often generated from both human and mouse samples. To evaluate CLUEY’s ability to estimate the number of cell types in independent datasets from both species, we tested CLUEY against alternative methods on four independent datasets from each species (Baron et al., 2016; Segerstolpe et al., 2016; Zeisel et al., 2015; Zilionis et al., 2019). These datasets were selected to ensure a varying number of cells and cell types between the two species, including the organs tested (Figure 4A). Our experiments on these datasets revealed that CLUEY significantly outperforms alternative methods on estimating true number of cell types especially in the Lung and the two Pancreas datasets (Figure 4B). In terms of cell clustering, CLUEY performed the best in the Lung dataset compared to other methods and show competitive results in the other three datasets (Figure 4C). It is worth noting that in some cases the number of cell types estimated by SC3 greatly deviate from the true number of cell types, resulting in poor clustering results (e.g. in the Lung dataset). Finally, compared to alternative methods, CLUEY has the advantage of offering interpretable clustering results, as it outputs both statistics and the most similar cell types in the knowledgebase used to guide the estimation process. This information can be utilised to further refine the clusters by guiding the annotation of clusters at a higher resolution, as well be demonstrated in the next section.

**Figure 4.**
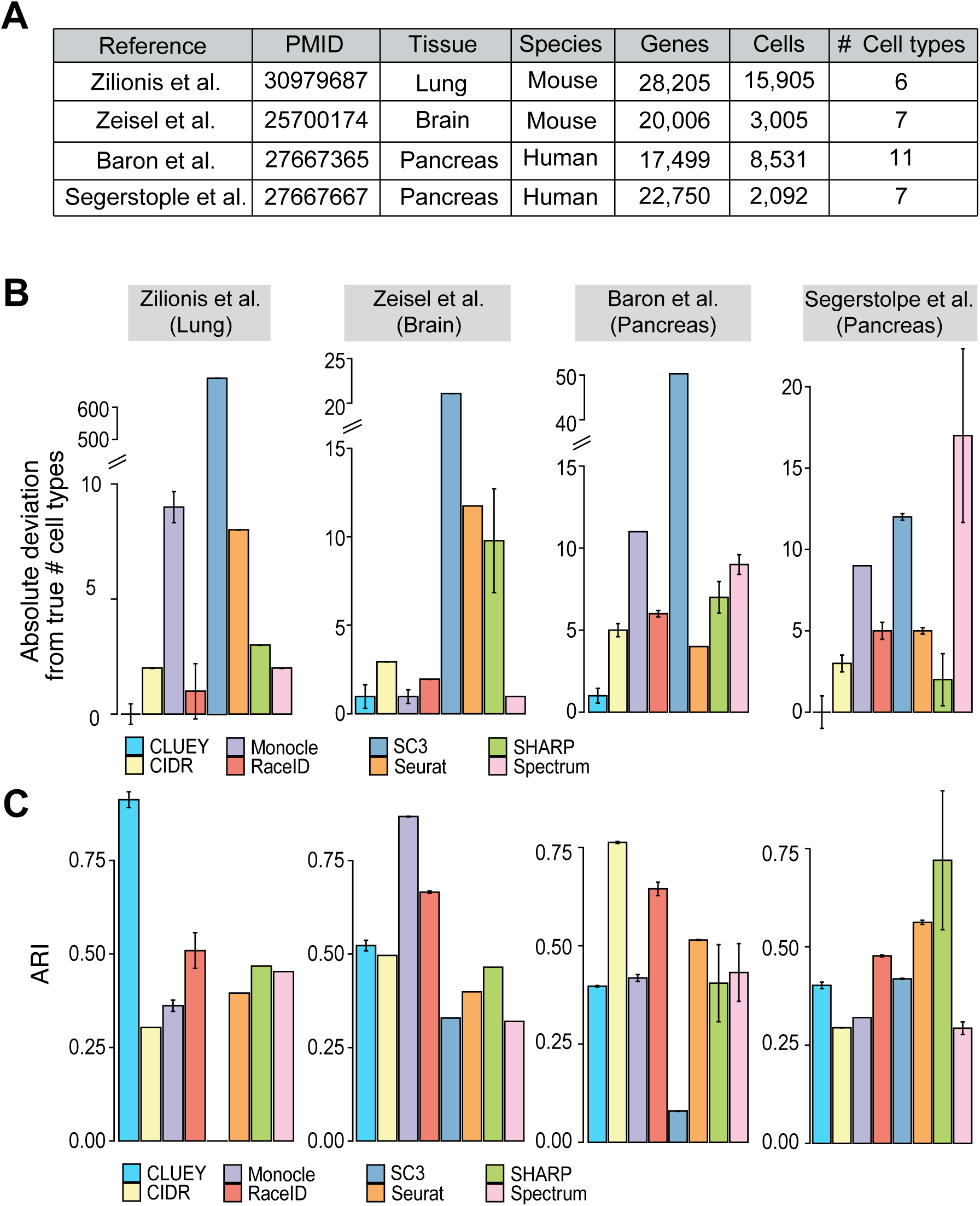
Performance on independent mouse and human datasets. **(A)** Description of datasets used for independent evaluation. **(B)** Absolute deviation on number of cell type estimation for each method on the four datasets. **(C)** For each clustering method, ARI quantification of cell clustering with respect to the original cell type labels from each study.

### CLUEY enables multimodal clustering without the need for a multi-omic knowledgebase

To assess CLUEY’s utility and performance in cluster single-cell multi-omics data, we compared it to Seurat across two datasets. These include the TEA-seq and CITE-seq datasets generated by Swansan et al. and Stephenson et al., respectively. In these experiments, we used the knowledgebase generated from HCA 10x data. Our results demonstrated high cluster concordance and accurate cell type annotation by CLUEY compared to the cell type labels from the original studies in both datasets (Figure 5). In comparison, Seurat appears to overestimate the number of cell types and yield lower ARI values compared to CLUEY (Figure 5B, 5D). Notably, while CLUEY also overpredicted the number of clusters in the TEA-seq dataset (Figure 5A), all clusters corresponded to immune cells, aligning closely with the original cell type labels. For instance, CLUEY grouped cells originally labeled as ‘Mono CD16’ in the TEA-seq data as a single cluster which correlated with the ‘Mature NK T Cell’ in the knowledgebase, and cells labeled as ‘Mono CD14’ were also grouped together and most correlated with ‘Dendritic Cell’ and ‘Classical Monocyte’, consistent with their known CD14 expression. Conversely, although CLUEY underestimated the number of cell types in the CITE-seq dataset, it accurately grouped cells into clusters that were informed by the related broader cell types. For example, cells labeled as ‘NK 16hi’ and ‘NK 56hi’ in the original data were correctly grouped as a cluster which correlated most with ‘NK Cell’ (i.e. Natural Killer cells), as the finer cell types were not present in the knowledgebase generated from the HCA 10X. These findings underscore CLUEY’s ability to cluster multimodal data without requiring a matched multimodal dataset, expanding its utility as multi-omic datasets become more prevalent.

**Figure 5.**
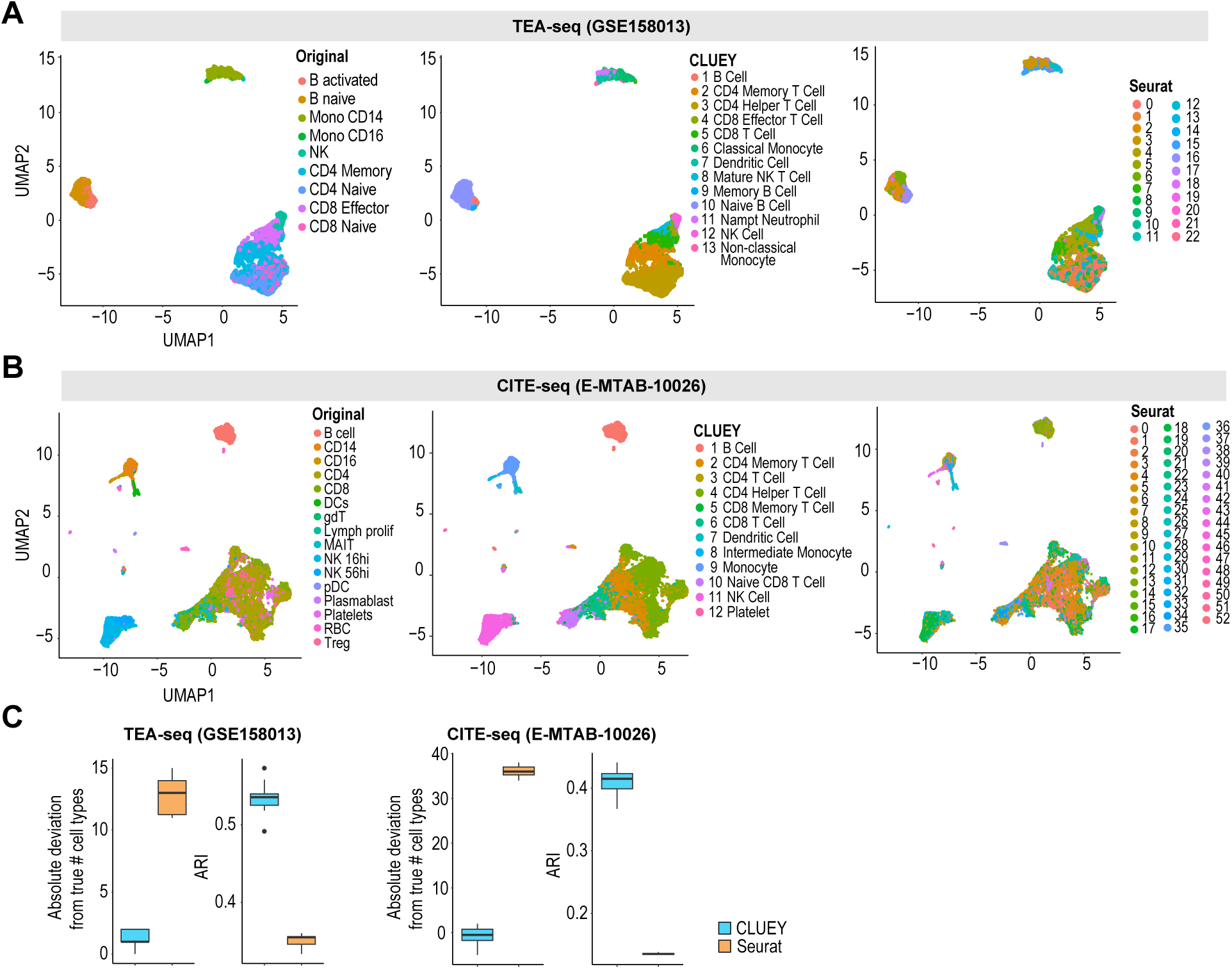
CLUEY enables multimodal clustering and annotation without the need for a multi-omic knowledgebase. **(A,B)** UMAPs of original cell type labels, closest matching cell types in the knowledgebase output by CLUEY, and Seurat cluster IDs, for (A) TEA-seq data, and (B) TEA-seq data. Both methods were supplied with RNA and ATAC modalities as input, and CLUEY utilised a knowledgebase generated from uni-modal HCA 10X data. **(C)** Boxplots of the Absolute Deviation and ARI between the original cell type labels and predictions of CLUEY and Seurat for TEA-seq and CITE-seq. Dataset was down-sampled to 90% of the original number of cells 10 times to capture variability.

## Discussion

In this work, we have presented CLUEY, a knowledge-guided framework for biologically interpretable clustering. CLUEY employs a novel recursive clustering algorithm and jointly utilizes differentially stable genes to determine the optimal number of clusters while providing the closest matching reference cell types for interpretability. Importantly, CLUEY averages the expression profiles of each cluster to minimize noise and capture more signal, thus enabling stable performance. In this study, we have demonstrated CLUEY’s performance in a variety of practical scenarios. CLUEY enables stable clustering results, while providing interpretable biological information.

Advances in sequencing technologies recently have enabled us to capture multiple layers of molecular information within individual cells. While single-cell transcriptomics pioneered entry into the single-cell era, it only captures one layer of cellular biology. Complementing transcriptomic data with additional modalities, such as chromatin accessibility by ATAC-seq and surface protein expression by CITE-seq may enhance analyses, including clustering and annotation (Tang et al., 2023; Yu et al., 2023). In light of this, CLUEY has the capacity to integrate multiple modalities to harness the information within multimodal data and detect the number of cell types using this information. CLUEY enables multimodal data clustering without requiring a corresponding multi-omic knowledgebase. Thus, users have the flexibility of providing their own gene sets using only RNA modality, derived from either uni- or integrated multimodal data, further enhancing its versatility and applicability. Nevertheless, identifying the optimal approach for integrating unmatched multimodal data remains a challenge, and multiple methods have been recently developed for achieving this task (Argelaguet et al., 2021; Lee et al., 2023). Future work will involve determining an efficient strategy for integrating unmatched multi-omic datasets.

Another challenge is accurately deciphering cell state dynamics. While conventional methods typically categorise cell type identity as discrete states, cells of the same cell type often exhibit heterogeneity (Trapnell, 2015). This continuous trajectory of cell states poses a challenge for determining the threshold for clustering cells as belonging to one cell type or the other. Like many methods, CLUEY treats cell type identities as discrete and groups a cell to a cluster which is most correlated to a cell type in the knowledgebase. However, CLUEY provides the correlation scores for each of the clusters, recognising that a cell’s identity is continuous and often will not perfectly match the average profile of its most correlated cell type in the knowledgebase. Additionally, CLUEY identifies and treats cell type identity markers as unique to the cell types in the knowledgebase. While this may be appropriate for capturing cell identity, changes in cell state are often due to minor perturbations in the regulatory networks that determine cell identity. Discrete categorization of cell types simplifies analysis and interpretation but can overlook more subtle molecular variations that occur as cells move from one cell state to another. Thus, it will be important to capture the inherent relationships between genes and how they change within cell states to better identify cell state changes which can further inform the clustering process and lead to more biologically interpretable and accurate clustering.

Beyond cell types and states, it is possible that cells in the query data are not present in the knowledgebase. Some methods define a minimum threshold to designate these cells as unassigned or potentially novel. Yet this parameter is often arbitrarily determined with minimal biological relevance (Aran et al., 2019; Kong et al., 2022). Given CLUEY is a knowledge-based clustering method, we have opted for the current approach to provide correlation scores associated with the final clusters to provide as much information for further downstream analyses. For example, potential absent or novel cell types can be inferred and further explored using the correlation scores and provided by CLUEY. While a threshold may be easier to interpret, we believe it can bias the conclusions drawn as the appropriate minimum threshold will change depending on many factors, including the sequencing platforms used to generate the data. This approach allows CLUEY to group cells together that are most correlated with a cell type in the knowledgebase but does not strictly assign them to a cell type. Importantly, the final clusters can have heterogeneous correlations scores, allowing users to identify potential novel subpopulations in the data.

## Conclusion

In summary, CLUEY is a method that enables knowledge-guided cell type detection and clustering of single-cell omics data. It facilitates the integration and clustering of multi-omic query data without the need for a matching multi-omic knowledgebase and performs a novel recursive clustering algorithm to generate stable and robust results. The knowledge-guided approach implemented in CLUEY marks a step towards biologically-informed clustering and for providing researchers with biologically interpretable clustering, thereby enhancing informed decision-making in subsequent downstream analyses.

## Data and code availability

All data used in this study are publicly available (see Material and Methods). Code for CLUEY is available at https://github.com/SydneyBioX/CLUEY.

## Acknowledgments

The authors thank their colleagues at the Sydney Precision Data Science Centre for their support and intellectual discussions. They would also like to thank Andy Tran, Alex Qin, Farhan Ameen, Sanghyun Kim and Hani Kim for their feedback and edits.

## Author Contribution

P.Y, conceived and funded the study. D.K, C.C completed the analysis and design of this study with the guidance from P.Y and J.Y.H.Y. The implementation and construction of the R package for the case study were done by D.K and L.Y. This article was completed jointly by all authors and all authors wrote, reviewed and approved the manuscript.

